# Salt corrections for RNA secondary structures in the ViennaRNA package

**DOI:** 10.1101/2023.04.07.536000

**Authors:** Hua-Ting Yao, Ronny Lorenz, Ivo L. Hofacker, Peter F. Stadler

## Abstract

**Background:** RNA features a highly negatively charged phosphate backbone that attracts a of cloud counter-ions that reduce the electrostatic repulsion in a concentration dependent manner. Ion concentrations thus have a large influence on folding and stability of RNA structures. Despite their well-documented effects, salt effects are not handled by currently available secondary stucture prediction algorithms. Combining Debye-Hückel potentials for line charges and Manning’s counter-ion condensation theory, Einert *et al*. [*Biophys. J*. **100**: 2745-2753 (2011)] modeled the energetic effects contributions monovalent cations on loops and helices.

**Results:** The model of Einert *et al*. is adapted to match the structure of the dynamic programming recursion of RNA secondary structure prediction algorithms. An empirical term describing the dependence salt dependence of the duplex initiation energy is added to improve co-folding predictions for two or more RNA strands. The slightly modified model is implemented in the `ViennaRNA` package in such way that only the energy parameters but not the algorithmic structure is affected. A comparison with data from the literature show that predicted free energies and melting temperatures are in reasonable agreement with experiments.

**Conclusion:** The new feature in the `ViennaRNA` package makes it possible to study effects of salt concentrations on RNA folding in a systematic manner. Strictly speaking, the model pertains only to mono-valent cations, and thus covers the most important parameter, i.e., the NaCl concentration. It remains a question for future research to what extent unspecific effects of bi- and tri-valent cations can be approximated in a similar manner.

**Availability:** Corrections for the concentration of monovalent cations are available in the `ViennaRNA` package starting from version 2.6.0.

## Introduction

Nucleic acids are highly negatively charged molecules since their phosphate backbone carries one negative charge per residue. Structure formation brings these charges into close proximity and thus incurs substantial electrostatic penalties on the secondary and tertiray structure level. Nucleic acid–ion interactions also provide large interaction energies and therefore contribute decisively to the folding RNA and DNA and to their interactions with ligands and macromolecule partners [1]. Counter-ions reduce the electrostatic repulsion of the backbone. Cation concentrations determine the extent of this “charge screening” and thus strongly influence RNA folding. Indeed, many functional RNAs will not fold under low salt conditions [2], and experimental investigations of the thermodynamics of RNA folding are mostly confined to high salt conditions. Energy parameters for RNA secondary predictions likewise pertain to 1M sodium concentrations, more precisely 1.021M while taking all Na^+^ ions in the buffer into account [3]. Although the importance of counter-ions for the RNA folding is well known, ion concentration, in contrast to temperature, is not a tunable parameter in currently available RNA secondary structure prediction tools. While temperature dependence is conceptually straightforward and can be easy modeled by splitting free energy contributions into enthalpic and entropic contributions [4], the energetics of ion–nucleic acid interactions are much more difficult to understand.

Cations affect RNA structure in two different ways. The electrostatic stabilization of the structure due to charge screening is at least conceptually independent of the chemical nature and charge of the cation. In addition, divalent cations and in particular Mg^2+^ can also strongly bind to specific, chelating sites [5, 6]. Quantitative salt effects on RNA folding have been studied extensively over the last decades, see [7] for a study that summarized much of the pertinent earlier literature. In the absence of a well-founded theoretical model, most authors resorted to describing the salt-dependence of RNA folding by means of simple heuristic function fitting the effects of changes in the sodium concentration on the free energy of folding or a melting temperature. Such empirical fits, however, are limited to handling salt effects close to standard conditions, and an approach that explains the functional form of salt effects is clearly preferrable.

If three-dimensional structures are known, the nonlinear Poisson-Boltzmann equation can be solved obtain electrostatic potentials of RNA molecules in solution [8–10]. This approach, however, appears to be too detailed to derive a practically manageable parametrization of salt effects at the level of secondary structure prediction algorithms. In order to handle counter-ions in RNA secondary structure prediction algorithms, the effects must be attributed to individual bases, base pairs, or loops (including the stacking of two consecutive base pairs). This is necessary because secondary structure prediction algorithms operate on these combinatorial substructures [11]. This, in particular, precludes models that explicitly require a detailed geometric description of three-dimensional structure of an RNA.

To derive a suitably simple model, Einert and Netz [12] proposed to represent the RNA backbone as a charged polymer that interacts by means of a Debye-Hückel potential [13] and treats single-stranded regions as freely jointed chains [14]. The non-linear screening effect of monovalent cations is incorporated using Manning’s approach to counter-ion condensation [15]. The formulation for loop contribution was orginally developed to understand the salt-dependent modulation of nucleosomal structures [16]. The two strands of a helix are modeled as parallel rods that again interact via a Debye-Hückel potential governed by the screening length. Here, the theory yields a per-position contribution that is independent of sequence features and the position within the helix [12]. While the theory makes several approximations it has been shown by its authors to reproduce experimental data quite well. It also has the advantage that it has no free parameters other than well known generic geometric characteristics of RNA 3D structures such as distances between nucleotides or the planes of stacked pairs. Of course, the secondary structure based approach has signficant limitations. In particular, it is by design not suitable to model the site-specific binding of chelated Mg^2+^, which in some cases is known to be crucial for tertiary structure formation and RNA function [17].

Here we implement in the `ViennaRNA` package [18] the Debye-Hückel/Manning model, which captures the energetic effects of electrostatic interactions of monovalent cations with structured RNA [12]. In the following theory section we briefly review the features of the energy model and show that it can be brought into a form where only the energy parameters but not the dynamic programming recursions are modified. We then evaluate the model on a collection of empirical data from the literature to show that use of this type of salt corrections has a significant beneficial effect.

### Theory

The model of [12] considers loops a freely jointed charged chains and helices as parallel rods interacting via Debye-Hückel potentials and uses Manning’s framework to model counter-ions condensation. This results in distinct types of correction terms from loops and helices, which we describe separately in the following. In either case we focus on how the salt correction terms are incorporated into the dynamic programming schemes for RNA secondary structure prediction. As we shall see, the salt corrections can phrased in such a way that they exclusively effect the energy parameters. The folding algorithms therefore remain unchanged. Owing to the architecture of the `ViennaRNA` package, it is therefore possible handle salt effects exclusively as a pre-processing the energy parameter set.

#### Salt corrections for loops

The electrostatic free energy contribution for a loop comprising *L* backbone bonds can be written, at the level of the Debye-Hückel approximation, in the form [16]:

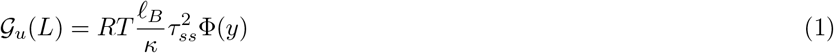

with *y* = *κl*_*ss*_*L* where *l*_*ss*_ = 6.4Å is the length of an RNA backbone bond, *ℓ*_*B*_ is the Bjerrum length and *κ*^−1^ is the Debye screening length, which depends on the ionic strength, and *τ*_*ss*_ = min(1*/l*_*ss*_, 1*/ℓ*_*B*_*z*_*c*_) accounts for the nonlinear electrostatic effects. For monovalent ions, the ionic strength equals the salt concentration *ρ* and thus we have *κ* = *κ*(*ρ*). Following [16], Φ is given by

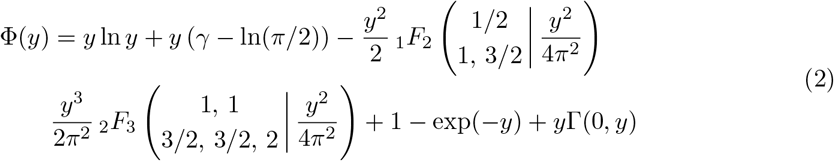

Here, _1_*F*_2_ and _2_*F*_3_ are generalized hypergeometric functions and Γ(0, *y*) is the incomplete gamma function. Using _1_*F*_2_(…, 0) = _2_*F*_3_(…, 0) = 1, and *y* ln *y* → 0 for *y* → 0, we observe that Φ(*y*) = 0. Equ.(A8) in [16] gives an exapansion for small *y* of form Φ(*y*)*/y* = (1 − ln(*π/*2)) + *O*(*y*^2^), where the ln *y* term and the logarithmic divergence of Γ(0, *y*) cancel. Thus, Φ(*y*) increases linearly with *y*. Since both *y* and *κ* are proportional to 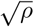, the *ρ*-dependence cancels and 𝒢_*u*_(*L*) approaches *L* times a constant for *ρ* → 0.

Since 𝒢_*u*_(*L*) describes the salt-dependent electrostatic effects on loops, this term is already included in the empirical energy parameters of the RNA standard model for the standard conditions of *T* = 37°*C* and 1*M* sodium concentration. Writing 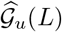 for the values on Eq.(1) at standard conditions, allows us to write the salt correction of a given loop as

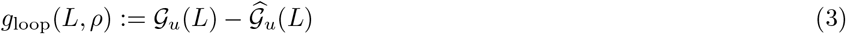

The `ViennaRNA` package quantifies the length of a loop by the number *m* of unpaired nucleotides rather than the number of bonds. For hairpin loops we have *L* = *m* + 1, while *L* = *m* + 2 for interior loops. In multi-loops we have *L* = *m* + *q* + 1, where *q* is the degree (number of branches) of the multiloop. Setting *q* = 0 for hairpin loops and *q* = 1 for interior loops, the appropriate salt correction for a loop is therefore *g*_loop_(*m* + *q* + 1, *ρ*).

As seen in Fig. 1, the salt correction, as expected, increases while decreasing the salt concentration contributing to the destabilization of the structure in a low salt environment. In the usual temperature range, the plot shows that loop correction is closed to a constance with a slight increase at low concentration.

**Figure 1.**
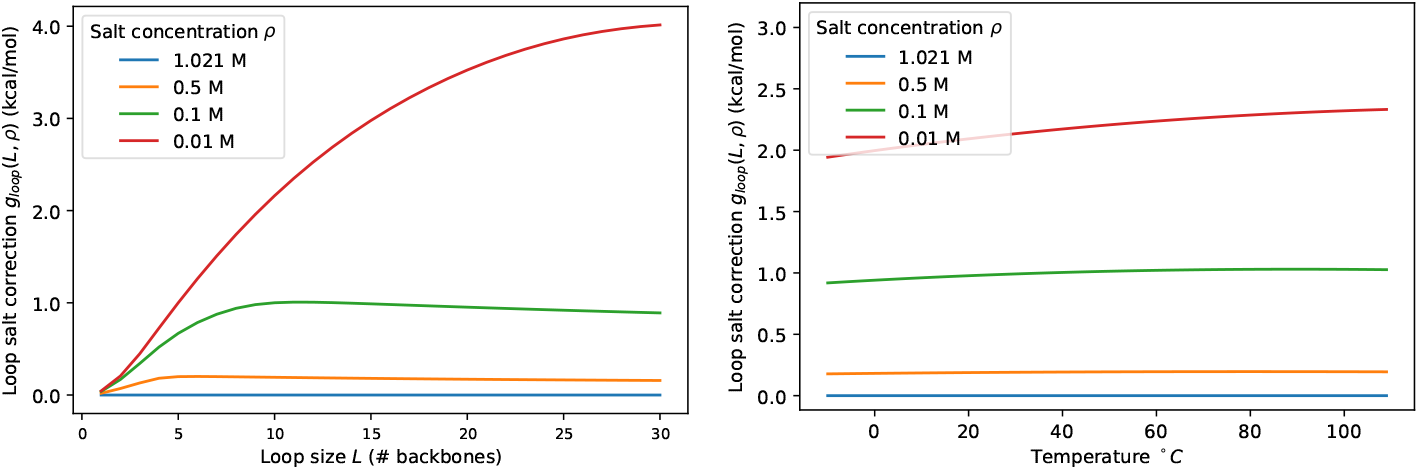
Loop salt correction *g*_loop_(*L, ρ*) as a function of loop size *L* = *m* + *q* + 1 (left) and as a function of temperature (right) for a fixed loop at size *L* = 10 for different salt concentrations *ρ*.

#### Linear approximation for multi-branch Loop

Although the conformation energy of a loop asymptotically depends on logarithmically on the length [19], this behavior is usually approximated for multi-loops by a linear function

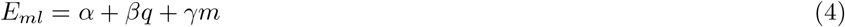

for the sake of computational efficiency. The parameters *α, β*, and *γ* are the energy cost for having, respectively, the closing pair, branch, and unpaired base in a multiloop. Without this approximation, the dynamic programming recursions of the folding algorithm become cubic in their memory consumption since loop lengths need to be tracked explicitly [20]. The linear approximation is further motivated by the empirical observation that models with logarithmic dependence are outperformed by the linear approximation in terms of accuracy of structure prediction [20]. In order to handle multi-loops without abandoning the memory-efficient multiloop decomposition, we retain the linear multiloop energies and employ a linear fit of the form

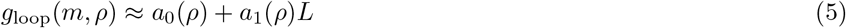

to model the salt-corrections. Again using *L* = *m* + *q* + 1, this amounts to adding *a*_0_+*a*_1_ to the closing pair term *α* of the multi-loop and *a*_1_ to both each unpaired base *γ* and each component*β* of the multi-loop. The salt-dependent multiloop models thus reads:

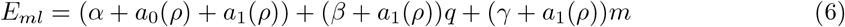

In order to fit the multi-loop parameters in practise, we first investigated their size distribution in a sample of 5000 MFE structures for different structure sizes, see Fig. S1. Although multi-loops with size *L* = 3 exist, very short multi-loops are rare. We argue that inaccuracies in these rare cases are likely acceptable. Very short loops presumable are also subject to specifically contrained three-dimensional structures and thus follow the model only to a crude approximation in the first place. In the current implementation, we use the loop size range *L* ∈ [6, 24] to obtain linear fits for *a*_0_ and *a*_1_ from *g*_loop_(*L, ρ*), see Fig. 2 for salt correction and their linear approximations. In general, the fit over-corrects for very small loops and, at low salt concentrations, also for very large loops. The maximal fitting errors are on the order of 1 kcal/mol, which is still within the ball-park of the rather large inaccuracies expected for multi-loop energies.

**Figure 2.**
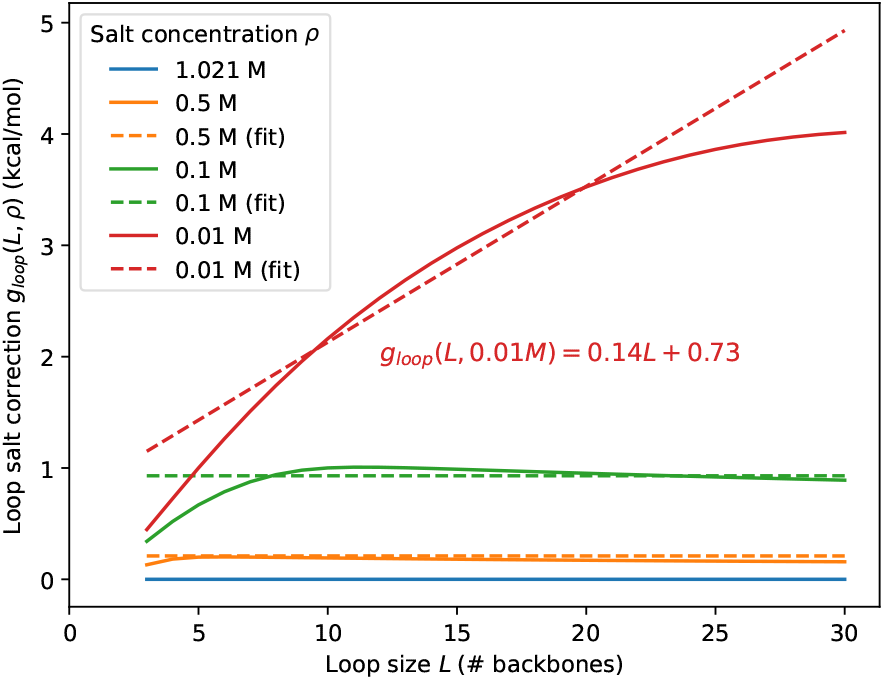
Loop salt correction (solid) and linear approximation (dashed) in the function of loop size *L* at different salt concentration.

#### Salt corrections for stacked base pairs

Describing the backbones of stacked pairs as rods with distance of *d* = 2nm interacting via a Debye-Hückel potential with screening length 1*/κ* yields the following electrostatic energy for the interaction of a nucleotide with the other strand [12]:

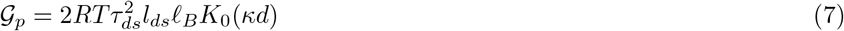

Here the charge density *τ*_*ds*_ = min(1*/l*_*ds*_, 1*/*(*ℓ*_*B*_*z*_*c*_)) is again estimated according to Manning’s counterion condensation theory [15]. The length parameter *l*_*ds*_ = 3.4Å is the helical rise per base pair and *z*_*c*_ = 1 is the charge of the cation. *K*_0_ denotes the zeroth-order modified Bessel function of the 2nd kind, see e.g. [21] and Fig. S2. Since *K*_0_(*z*) diverges like −ln *z* for *z* → 0, and 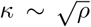, the salt corrections 𝒢_*p*_ diverges logarithmically for vanishing salt concentrations.

As in the case of loops, these electrostatic effects are already included in the empirical energy parameters for standard conditions. The relevant salt correction thus is given by the difference between the values 𝒢_*p*_ at the current conditions and standard conditions.

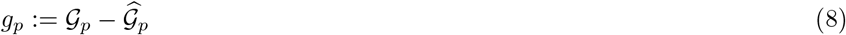

Fig. 3 shows the dependence of the position-wise salt correction for stacking energies as function of salt concentration and temperature. Similar to the loop correction, the stack correction is closed to a constance in the usual temperature range for a given salt concentration.

**Figure 3.**
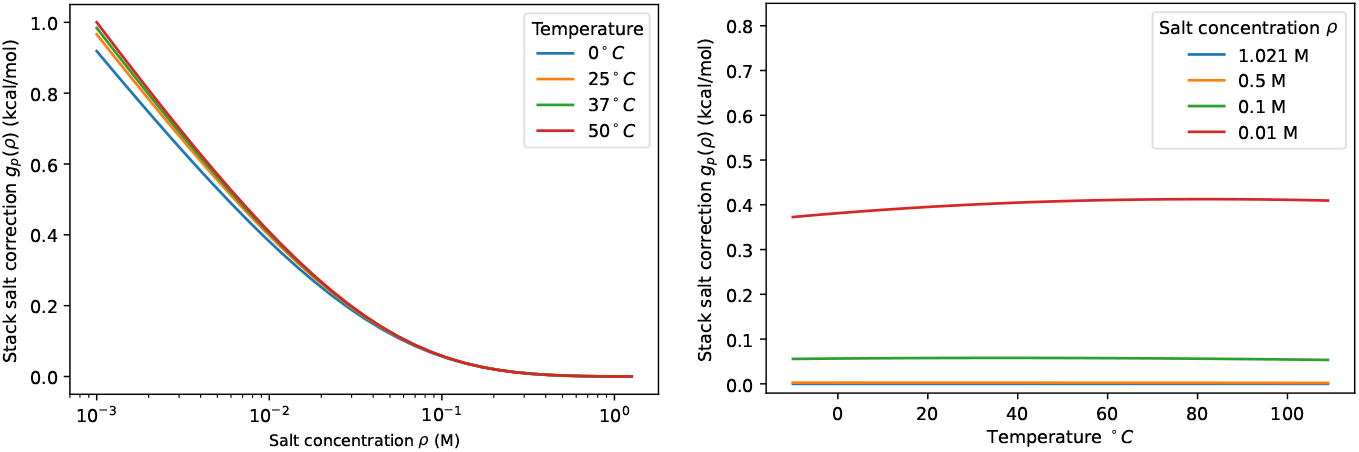
Salt correction for a stacked pair as a function of salt concentration (left) and temperature (right).

#### Salt corrections for duplex initialization

The formation of a double strand from two RNA molecules in solution is associated with an additional initialization energy *E*_init_ in the Turner energy model. One expects that duplex formation becomes more difficult due to electrostatic repulsion at low salt concentrations. The initialization energy thus should also depend on *ρ*. In addition, the distance between two single strands changes during formation, which was neglected in the theory as discussed in [12]. Indeed, as we will see later in the result section, the duplex free energies are systematically overestimated compared to the experimental data.

Due to the lack of theoretical support, we propose a salt corrections for duplex initialization *g*_init_(*ρ*) derived from the prediction and the experimental data taken from [7]. Let *g*_*w*_(*ρ*) and 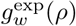 be the predicted and experimental salt correction at concentration *ρ* from the standard condition for a give duplex *w*. Fitting the difference 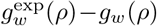 of 18 duplexes at four non-standard sodium concentrations yields, as plotted in Fig. 4, the salt corrections for duplex initialization

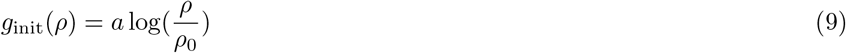

with *a* = −100.14 kcal/mol.

**Figure 4.**
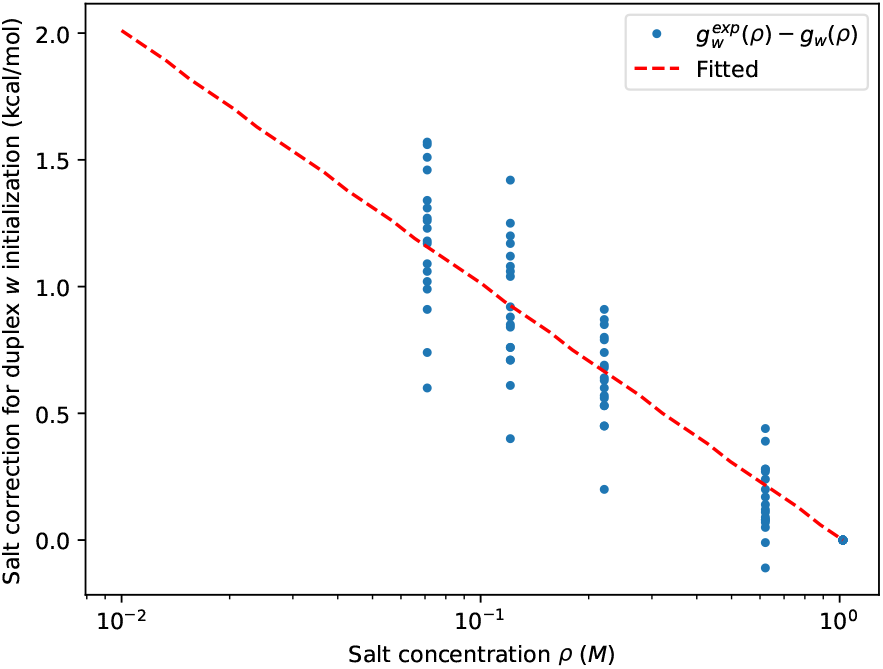
Salt correction for duplex initialization fitted (red) from the difference between experimental and predicted duplex salt correction (blue).

### Implementation in ViennaRNA

The extension of `ViennaRNA` provides access to four user-defined parameters: the concentation of the monovalent cation *ρ*, the bounds *L*_1_ and *L*_2_ delimiting the interval of loop length that is used to fit the two multiploop parameters *a*_0_ and *a*_1_ for given salt concentration, and the user-provided salt correction *g* for duplex initialization. Default values are *L*_1_ = 6, *L*_2_ = 24, salt concentration *ρ* = 1.021 M, and *g* = 0. In `ViennaRNA`, these parameters are appended in the model object vrna_md_s as salt for *ρ*, saltMLLower and saltMLUpper for *L*_1_ and *L*_2_, and saltDPXInit for *g*. For the salt concentration, `ViennaRNA` assumes the standard conditions of the Turner energy model, *i*.*e*., *ρ* = 1.021mol/l. Thus no salt corrections apply by default. If a different concentration *ρ* is requested, first the value of *g*_p_(*ρ*) for the stack and the array of values *g*_loop_(*L, ρ*) for loop of different size *L* ∈ [1, 31] is computed. Note that these energy contribution depend on the temperature *T* and thus are recomputed if the user sets a different temperature. In addition, the use of duplex initialization salt correction *g*_init_(*ρ*) for duplex is turned off if users set saltDPXInit to zero.

The array *g*_loop_ is then used to determine *a*_0_(*ρ*) and *a*_1_(*ρ*) by linear regression. Subsequently, the free energy parameters are set as sums of the default values *E*(0) for the given temperature and the salt corrections:

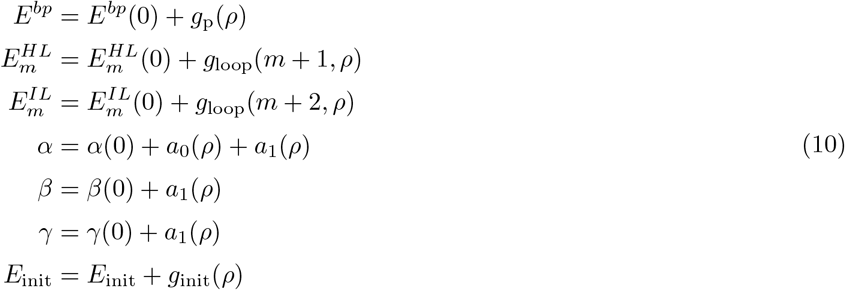

Here *E*^*bp*^, 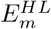, 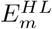, and *E*_init_ refer to *all* parameters for stacked pairs, hairpin loops of length *m*, interior loops of length *m*, and duplex initialization, respectively. Dangling end and coaxial stacking contributions, on the other hand, remain unchanged.

The Bessel functions *K*_0_ is computed as in `scipy`, which in turn used the `cephes` mathematical function library described in [22]. The function Φ in loop salt correction can be approximated as given in [12]:

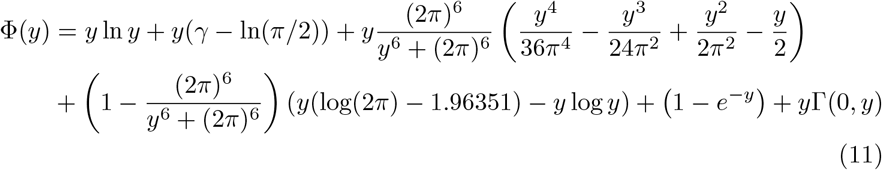

#### Parameters

The key physical parameter containing the salt correction terms are the Debye screening length *κ*^−1^. It is convenient to express *κ*^−1^ in terms of Bjerrum length *∓*_*B*_ and the ionic strength *I*:

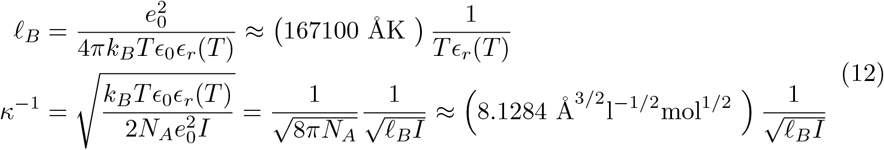

The constants appearing in these expressions are the unit charge *e*_0_ ≈ 1.602*·*10^−19^C, Avogadro’s constant *N*_*A*_ ≈ 6.022 *·* 10^23^mol^−1^, and the vacuum permissivity *ϵ*_0_ ≈ 8.854 10^−22^C^2^/JÅ. The salt concentration enters only through the ionic strength 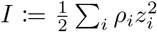, where *ρ*_*i*_ is the concentration and *z*_*i*_ is the charge of ion-species *i*. For monovalent ions, as in the case of NaCl, the ionic strength reduces to the salt concentration, i.e., *I* = *ρ*. The salt concentration *ρ* is expressed as molarity, i.e., in units of mol*/*l, whereas the Debye screening length *κ*^−1^ and the Bjerrum length *ℓ*_*B*_ are conveniently expressed in Å. Note that the conversion factor for the length units (l=dm^3^ versus Å) is absorbed into the numerical constant.

The temperature dependence of *E*_*r*_ can be fitted from empirical data. In the current implementation we use the function proposed in [23]:

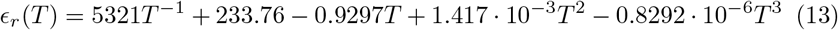

with temperature in Kelvin. The temperature-dependence of *ϵ* _*r*_ ensures that the Bjerrum length *ℓ*_*B*_ is longer than the backbone bond length *l*_*ss*_ in the entire temperature range, see Fig. S3. The nonlinear electrostatic effects on unpaired bases thus become *τ*_*ss*_ = 1*/*(*–*_*B*_) for the monovalent cations considered here.

### Comparison with experimental data

#### Available data sets

Even though the dependence of RNA structures on salt concentrations is of considerable practical interest, systematic data sets suitable for benchmarking salt correction models are by no means abundant. Most of the direct evidence for the salt dependence derives from studies of short duplexes and hairpins. Here, we analyzed the following three datasets:

1. Melting experiment of 18 RNA self-complementary duplexes of length 6 at the 8 different sodium concentration, 0.071, 0.121, 0.221, 0.621, and 1.021M, with different species concentration *c* were reported in [7, 24]. Both melting temperatures *T*_*m*_ and free energies at 37°*C* were are obtained from 1*/T*_*m*_ vs. ln *c* plots. The same data were also used by the same lab [24] to obtain optimised thermodynamic parameters, entropy and enthalpy, at a given salt concentration.
2. A different set of 8 self-complementary duplexes of length 10, 12, and 14 was reported in [25]. The melting temperature *T*_*m*_ is derived from the melting curve as the temperature at which half of the dimer dissolved. The data set covers two different species concentration *c*, 100*μM* and 2*μM* and two sodium concentrations, 1.0002*M* and 0.0102*M*.
3. The free energies of forming hairpins at different sodium concentrations were measured in [26]. The thermodynamics parameters at different salt concentration, different sodium concentration with an additional of 0.011*M* cations in the buffer, were obtained from melting experiments. Hairpins consist of a helix of length 5 and a hairpin loop of size 8 or 10.

#### Comparison of duplex free energies

Free energies were computed using `RNAcofold` [28, 29] with a self-complementary correction of *RT* ln 2 added for all sequences that coincided with their reverse complement. Since the predicted free energy at the standard condition differs from the experimental values, we are interested in comparing the salt correction *g*_*w*_(*ρ*) = *E*_*w*_(*ρ*) − *E*_*w*_(*ρ*_0_) at concentration *ρ* from the standard concentration *ρ*_0_ = 1.021M, where *E*_*w*_(*ρ*) is the free energy of duplex *w* at concentration *ρ*. Let *l*_*w*_ and gc_*w*_ be the length and the fraction of GC of duplex *w*. Then the salt correction for `RNAcofold` is then given by.

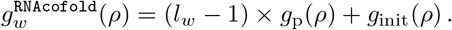

The salt correction proposed by Chen & Znosko [7] is

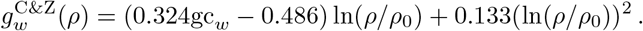

The computational results are summarized in Fig. 5. Not surprisingly, the Chen & Znosko salt correction provides a slightly better fit to the data because the empirical formula for 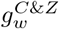 was obtained by fitting to the same data set. In contrast, only the duplex initialization *g*_init_(*ρ*) is fitted in our model. The largest deviations are observed for GC-only sequences.

**Figure 5.**
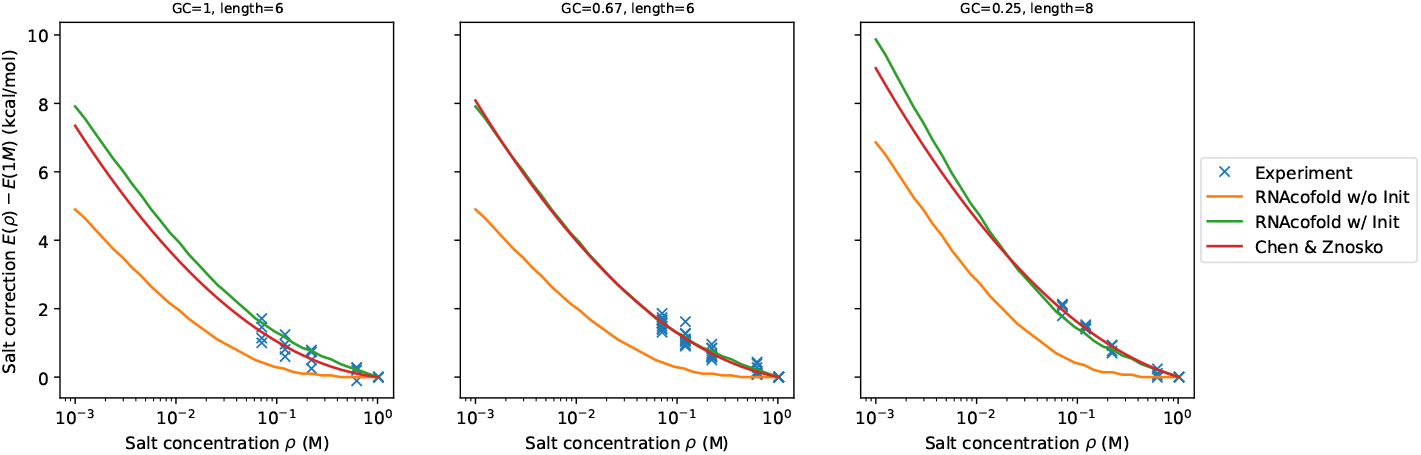
Duplex salt correction at different salt concentration from experiment (blue), Chen & Znosko (red), and `ViennaRNA` prediction with (green) and without (orange) salt correction for duplex initialization. The experimental free energy is derived from van’t Hoff plots of 1*/T*_*m*_ versus ln *c*. Most of the experimental values at 1M are taken from [27].

#### Comparison of duplex melting temperatures

Let *A* be a self-complementary RNA sequence and *AA* the corresponding dimer. The dimerization reaction is 2*A ⇌ AA*. The corresponding concentrations are denoted by [*A*] and [*AA*], respectively. In equilibrium, we have

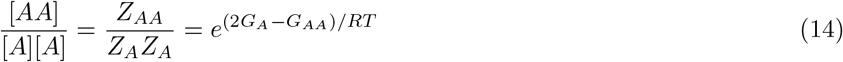

where *Z*_*A*_ and *Z*_*AA*_ are the partition functions of the monomer and the dimer, respectively. Here *G*_*A*_ = −*RT* ln *Z*_*A*_ and *G*_*AA*_ = −*RT* ln *Z*_*AA*_ are the *ensemble* free energies of *A* and *AA*, respectively. Note that in a pure two-state system, we can replace *G*_*A*_ and *G*_*AA*_ by the corresponding minimum free energies *E*_*A*_ and *E*_*AA*_, respectively. We define the melting temperature *T*_*m*_ as the temperature at which half of *A* forms the dimer *AA*, i.e., where [*AA*] = *c/*4 and [*A*] = *c/*2. Equ.(14) then yields

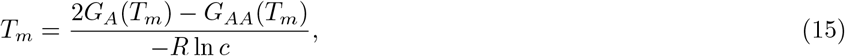

where we write *G*_*A*_(*T*_*m*_) and *G*_*AA*_(*T*_*m*_) to emphasize the temperature dependence of the ensemble free energies. The ensemble free energy *G*_*AA*_ for the pure dimer state is accessible directly via the function fc.dimer pf() within the `ViennaRNA` library. The correction for self-complementary is already taken into account during the computation of the the partition function [29]. Since the ensemble free energy is also a function of temperature, we use a binary search to find the melting temperature *T*_*m*_.

For the data set comprising 18 short duplexes, experimental melting temperatures are available for several distinct species concentrations *c*. Van’t Hoff’s equation

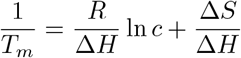

implies a linear relationship between changes in 1*/T*_*m*_ and changes in RNA concentration. We compare the predicted and experimental “van’t Hoff” plots, i.e., diagrams of 1*/T*_*m*_ *versus* ln *c*, see Fig. S4. Overall, we observe an excellent agreement on the slope between `RNAcofold` prediction and experiment. Predicted and experimental intercepts are slightly more different for few of the duplexes.

For the second dataset consisting of longer duplexes, we are interested in melting temperature correction Δ*T*_*m*_(0.01*M*) from the standard condition 1*M* since the data is only available for two species concentrations. Chen & Znosko [7] proposed the following fit

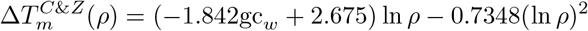

where gc_*w*_ is the GC fraction of duplex *w*. Fig. 6 shows the experimental melting temperature correction compared with 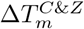 and the one computed by `RNAcofold`. Overall, the salt corrections described above show similar agreement (Pearson correlation *r* = 0.329) with the experiment as the empirical fit (*r* = 0.341) from [7].

**Figure 6.**
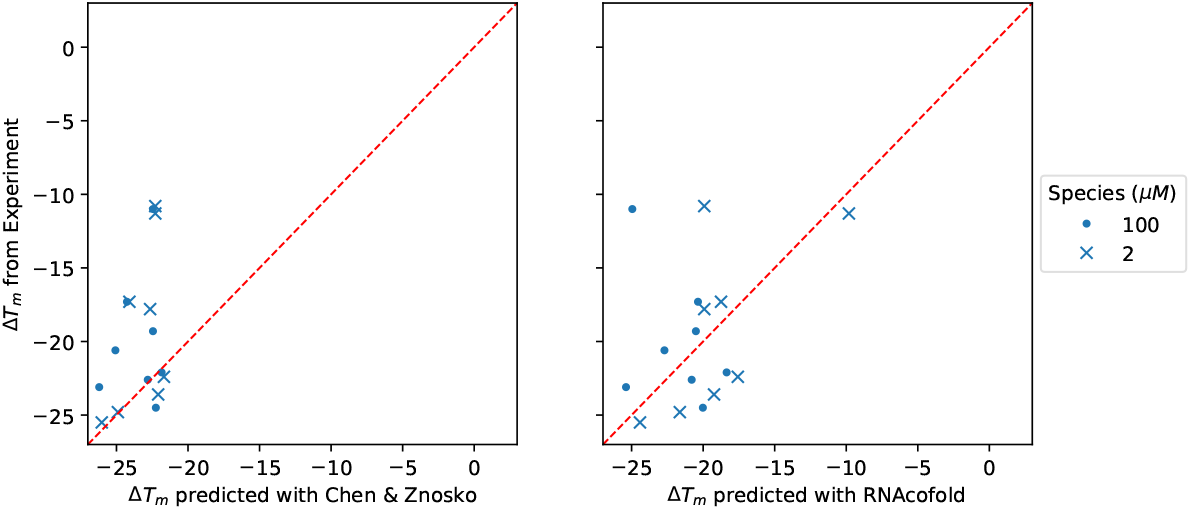
Comparison of experimental and predicted melting temperature corrections Δ*T*_*m*_ using the empirical fit by Chen & Znosko [7] (left) and `RNAcofold` with the salt corrections terms described in the present contribution (right)

#### Comparison of hairpin free energies

For the hairpins L8 and L10 described in [26], entropy, enthalpy, and free energy were derived from melting experiments. For comparison, we computed the free energy using `RNAfold` at 37°*C* for different salt concentrations *ρ*. Fig. 7 summarizes the experimental and predicted free energie corrections from 0.211 M. They are in very good agreement in particular for lower salt concentrations.

**Figure 7.**
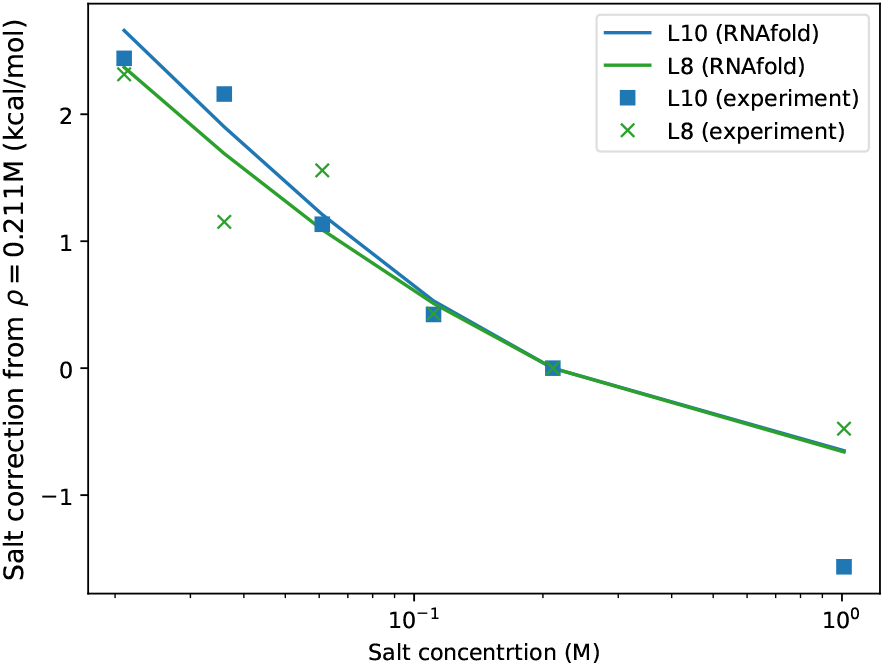
Comparison of experiment free energy correction of hairpin L8 (green) and L10 (blue) with RNAfold prediction.

## Discussion

Salt concentrations significantly influence folding and thermodynamics of nucleic acids. In this contribution we report on the implementation of an approximate physical model proposed by Einert and Netz [12] that represent the RNA backbone as a charged polymer interacting by means of a Debye-Hückel potential. The model was adapted to preserve the linear multi-loop model required for computational efficiency and extended by a empirical initiation term for duplex formation. While not perfect, the model reproduces experimental thermodynamic data on the NaCl dependence of folding energies and melting temperatures with reasonable accuracy. We note that the salt-dependent energies for stacks diverge logarithmically for vanishing salt concentrations *ρ*. The model thus cannot be used if cations are virtually absent.

The approach taken here has the practical advantage that it does not require any changes in the folding algorithms. The modification of the energy parameters is sufficient. The `ViennaRNA` packages handles this step during preprocessing. As a consequence, the compuational performance of the folding routines remain unchanged. Moreover, the salt corrections are consistently applied to all variants of the folding algorithms, e.g. minimum energy and partition function computations, consensus computations from alignments, and co-folding of two or more components.

Changes in salt concentration also affect the predicted secondary structures, see Fig. 8 for an example. To our knowledge, there are no detailed experimental data that document structural changes as a function of NaCl concentration so that a directed validation of structures predicted for low salt concentrations remains a topic for future research.

**Figure 8.**
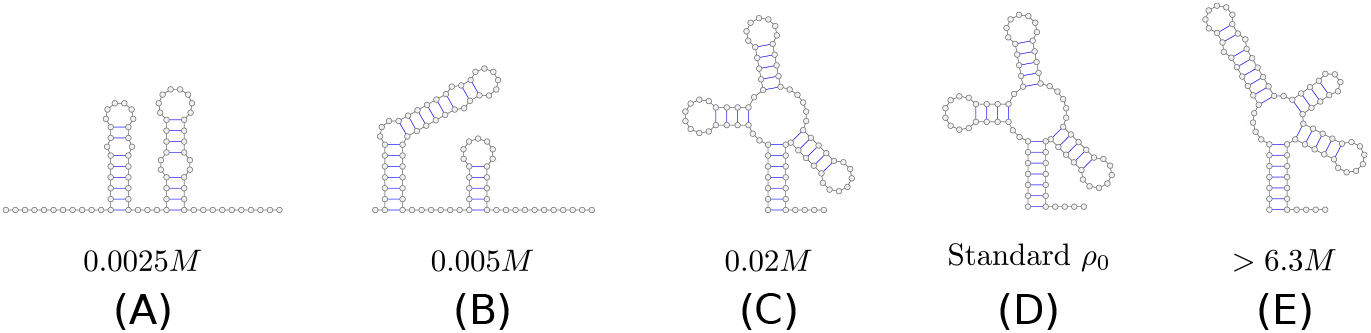
Examples of structural transitions. MFE structures of a tRNA sequence at different salt concentrations are predicted with RNAfold. Within the concentration range from 0.02 to 6.3M, the MFE structure is same as the one at the standard condition. The denaturation is observed at low concentration (A, B), while at high concentration (*>* 6.3*M*, corresponding to a saturated saline solution), both D and E can be the MFE. The tRNA sequence used is GCGGAUUUAGCUCAGUUGGGAGAGCGCCAGACUGAAGAUCUGGAGGUCCUGUGUUCGAUCCACAGAAUUCGCACCA.

The proposed salt correction diverges when the salt concentration becomes close to 0. The same behavior is also observed in our linear fitted correction for duplex initialization. To address it, we fitted a correction function with one parameter more comparing to Equ.(9) that converges at low concentration. Fig. S5 plots the predicted melting temperature correction versus the experiments of longer duplex, which shows a better agreement with Pearson correction *r* = 0.54. Due to the lack of data to validate the hypothesis of convergence at low concentration, we propose the simpler Equ.(9) as the main correction for duplex initialization.

In the current model, a basepair mismatch in a helix is treated as an 1*×*1 interior loop and thus is associated with the salt correction for loops at non-standard salt concentrations. However, such a mismatch is likely to result in a slighlty distorted helix that could still be seen as two parallel charged rods. One could therefore argue, that the salt correction for 2 stacked pairs rather than for a loop should be applied. Fig. 9 shows the difference of these two cases as a function of salt concentration. To our knowledge, there are no experimental data that could be used to decide which approach is more appropriate.

**Figure 9.**
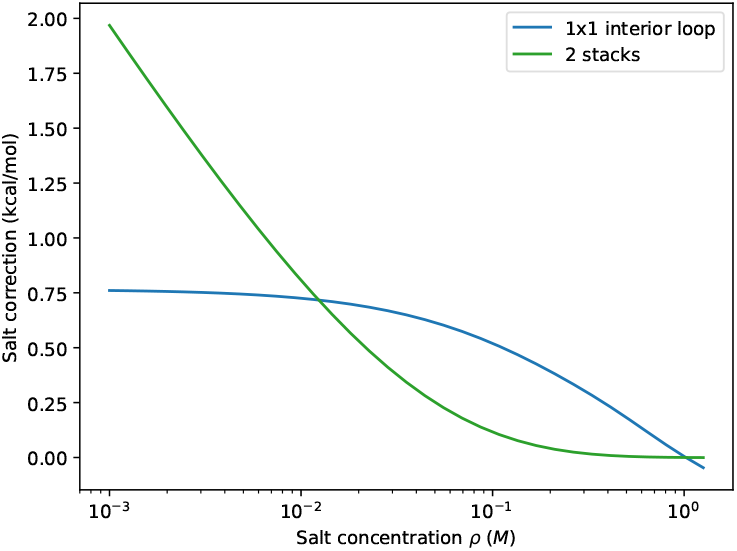
Free energy correction of an 1*×*1 interior loop (blue) and a helix of two stackings (green) at different salt concentration.

The approach taken here does not account for all effects of ions on RNA folding. Most importantly, it covers only unspecific interactions and thus does not describe specific interactions e.g. of Mg^2+^ with specific binding sites. Even for unspecific interactions, the validity of the model can be argued stringently only for monovalent ions [12]. At present, available data, e.g. [30], are not sufficient to test whether replacing *ρ* by the ionic strength is sufficient to reasonably account for mixtures of mono-valent and di-valent cations.

The model described here does not account for salt effects of RNA structures that are neither loops nor stacked base pairs. In particular it does not apply to G quadruplexes [31], which optionally can be included in secondary structure predictions [32]. Separate models for the ion dependencies of such features will need to be derived that account e.g. for the *K*^+^-dependent stabilization of RNA quadruplexes.

## Acknowledgements

This research was funded in part by the Austrian Science Fund (FWF), grant no. I 4520 and F 80, as well as the German Research Foundation (DFG), grant no. STA 850/48-1.

## Availability of data and materials

The implementation is available in the ViennaRNA Package starting with version 2.6.0 available at https://www.tbi.univie.ac.at/RNA.

## Ethics approval and consent to participate

Not applicable.

## Consent for publication

Not applicable.

## Authors’ contributions

All authors contributed to the design of the study and the writing of the manuscript. HTY implemented the model and conducted the computational evaluation, RL integrated the new features into the ViennaRNA package.

## Additional Files

Additional File 1 — List of supplementary figures

Figure S1 — Length distribution of multiloops

Distribution of multiloop size *L*, number of backbones, among MFE structures of 5 000 uniformly selected sequences at varied length.

Figure S2 — Approximation Error for *K*_0_

In [12] an approximation for the difference of *K*_0_ at a given concentration and 1*M* was proposed. However, we noticed that this approximation yields a non-vanishing salt correction at 1*M*. We therefore used the Cephes library to compute *K*_0_ directly. The panel shows the salt correction of base pair stack at 37*°C* in the function of salt concentration using the approximation (blue) and the precise computation implemented in ViennaRNA (orange).

Figure S3 — Nonlinear electrostatic effects *τ*_*ss*_

In [12], the permittivity (relative dielectric constant) *ϵ* _*r*_ of water *ϵ* _*r*_ ≈ 80 is assumed to be temperature independent. This assumption results in a discontinuity of *τ*_*ss*_ at around 53.3 *°C*. Incorporating the empirical temperature dependence of *ϵ* _*r*_ in equ.(13) [23] results in 1*/ℓ*_*B*_ *<* 1*/l*_*ss*_.

Figure S4 — Van t’Hoff plots for 18 duplexes.

Plotting 1*/T*_*m*_ versus ln *c* shows a generally good agreement of between predictions and the experimental data from from [7].

Figure S5 — Converged salt correction for duplex initialization.

Converged correction function fitted (left) to the difference 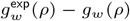 of 18 duplexes data [7], The plot (right) of the predicted melting temperature correction versus the experiments of longer duplexes [25] shows a better agreement with Pearson correction *r* = 0.54.

