## Additional File 1 for "Salt corrections for RNA secondary structures in the ViennaRNA package"

### Additional Files

Hua-Ting Yao, Ronny Lorenz, Ivo L. Hofacker, Peter F. Stadler

Supplementary Data

#### Additional file 1 — Length distribution of multiloops

Distribution of multiloop size $L$, number of backbones, among MFE structures of 5 000 uniformly selected sequences at varied length.

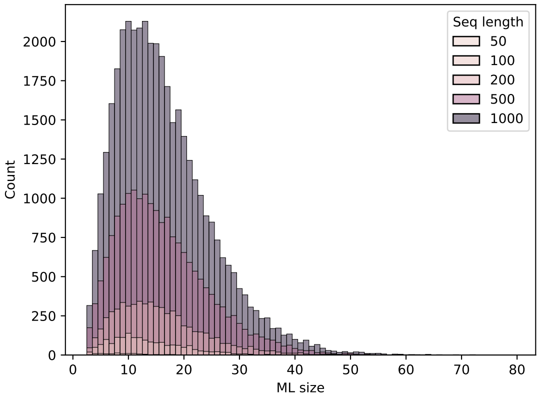

#### Additional file 2 — Approximation Error for $K_0$

In [1] an approximation for the difference of $K_0$ at a given concentration and $1M$ was proposed. However, we noticed that this approximation yields a non-vanishing salt correction at $1M$. We therefore used the Cephes library to compute $K_0$ directly. The panel shows the salt correction of base pair stack at $37°C$ in the function of salt concentration using the approximation (blue) and the precise computation implemented in `ViennaRNA` (orange).

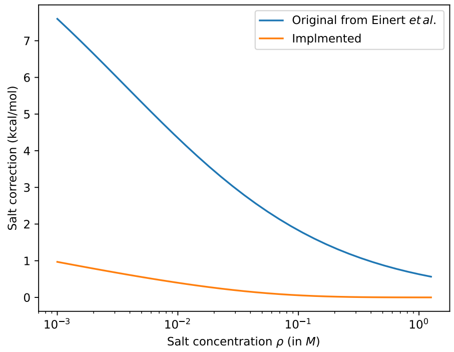

#### Additional file 3 — Nonlinear electrostatic effects $\tau_{ss}$

In [1], the permittivity (relative dielectric constant) $\epsilon_r$ of water $\epsilon_r \approx 80$ is assumed to be temperature independent. This assumption results in a discontinuity of $\tau_{ss}$ at around $53.3\,^{\circ}C$. Incorporating the empirical temperature dependence of $\epsilon_r$ results in $1/\ell_B < 1/l_{ss}$.

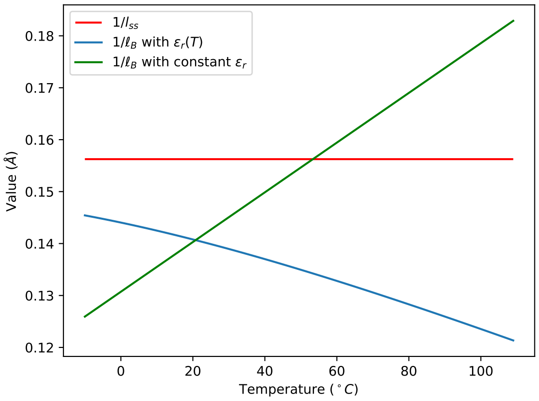

### Additional file 4 — Van t'Hoff plots for 18 duplexes.

Plotting $1/T_m$ versus $\ln c$ shows a generally good agreement of between predictions and the experimental data from from [2].

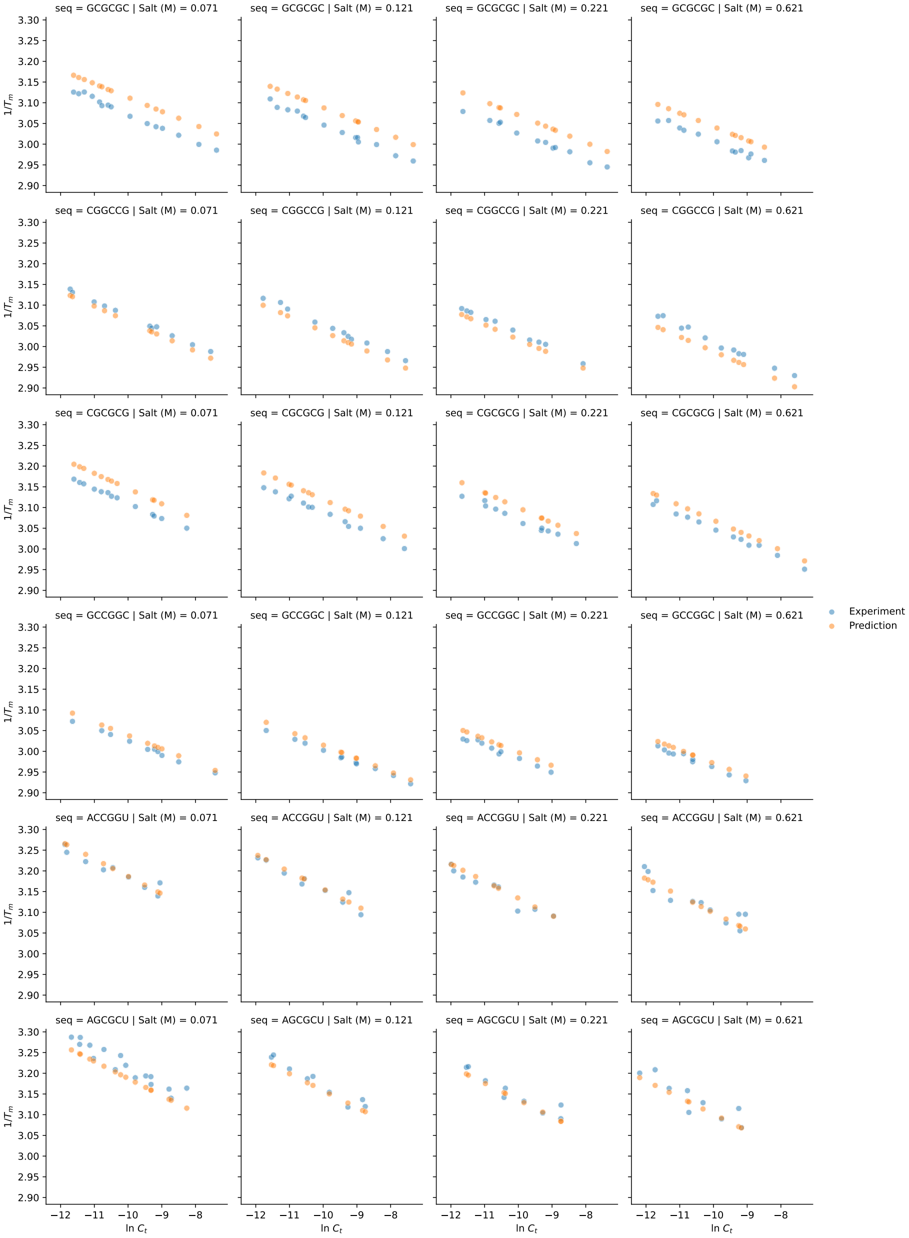

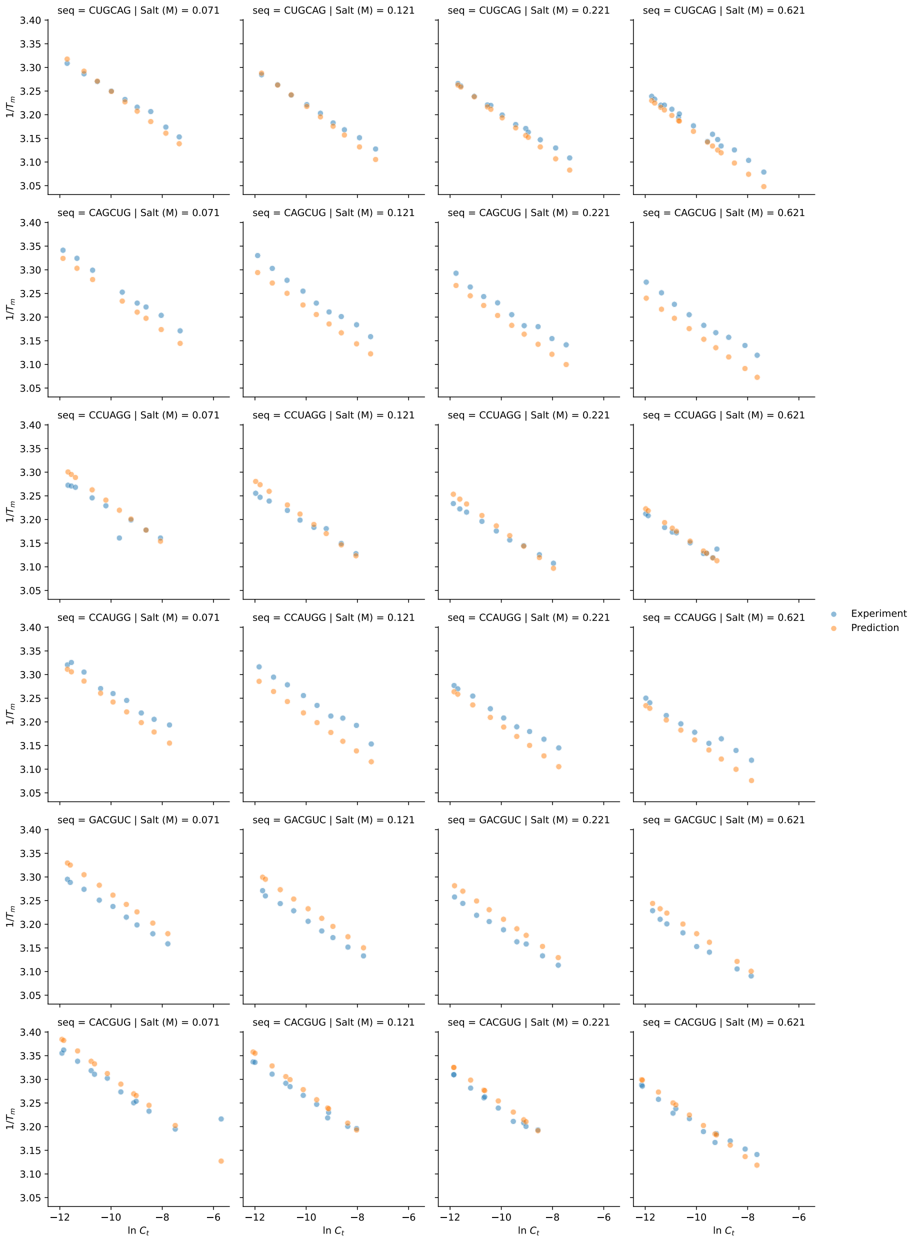
seq = CUGCAG | Salt (M) = 0.071
seq = CUGCAG | Salt (M) = 0.121
seq = CUGCAG | Salt (M) = 0.221
seq = CUGCAG | Salt (M) = 0.621
seq = CAGCUG | Salt (M) = 0.071
seq = CAGCUG | Salt (M) = 0.121
seq = CAGCUG | Salt (M) = 0.221
seq = CAGCUG | Salt (M) = 0.621
seq = CCUAGG | Salt (M) = 0.071
seq = CCUAGG | Salt (M) = 0.121
seq = CCUAGG | Salt (M) = 0.221
seq = CCUAGG | Salt (M) = 0.621
seq = CCAUGG | Salt (M) = 0.071
seq = CCAUGG | Salt (M) = 0.121
seq = CCAUGG | Salt (M) = 0.221
seq = CCAUGG | Salt (M) = 0.621
seq = GACGUC | Salt (M) = 0.071
seq = GACGUC | Salt (M) = 0.121
seq = GACGUC | Salt (M) = 0.221
seq = GACGUC | Salt (M) = 0.621
seq = CACGUG | Salt (M) = 0.071
seq = CACGUG | Salt (M) = 0.121
seq = CACGUG | Salt (M) = 0.221
seq = CACGUG | Salt (M) = 0.621
$1/T_m$
ln $C_t$
Experiment
Prediction

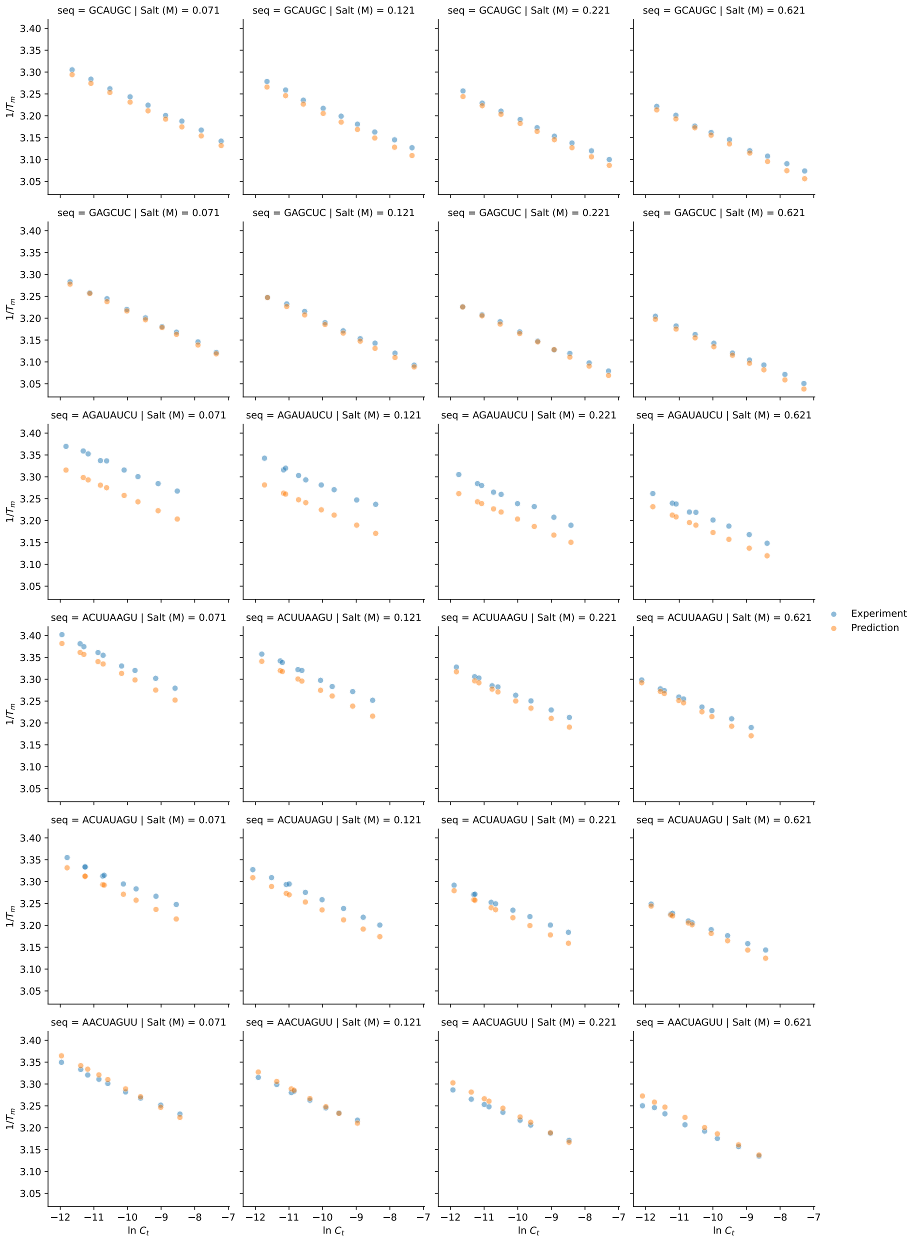
seq = GCAUGC | Salt (M) = 0.071
seq = GCAUGC | Salt (M) = 0.121
seq = GCAUGC | Salt (M) = 0.221
seq = GCAUGC | Salt (M) = 0.621
seq = GAGCUC | Salt (M) = 0.071
seq = GAGCUC | Salt (M) = 0.121
seq = GAGCUC | Salt (M) = 0.221
seq = GAGCUC | Salt (M) = 0.621
seq = AGAUAUCU | Salt (M) = 0.071
seq = AGAUAUCU | Salt (M) = 0.121
seq = AGAUAUCU | Salt (M) = 0.221
seq = AGAUAUCU | Salt (M) = 0.621
seq = ACUUAAGU | Salt (M) = 0.071
seq = ACUUAAGU | Salt (M) = 0.121
seq = ACUUAAGU | Salt (M) = 0.221
seq = ACUUAAGU | Salt (M) = 0.621
seq = ACUAUAGU | Salt (M) = 0.071
seq = ACUAUAGU | Salt (M) = 0.121
seq = ACUAUAGU | Salt (M) = 0.221
seq = ACUAUAGU | Salt (M) = 0.621
seq = AACUAGUU | Salt (M) = 0.071
seq = AACUAGUU | Salt (M) = 0.121
seq = AACUAGUU | Salt (M) = 0.221
seq = AACUAGUU | Salt (M) = 0.621
$1/T_m$
$\ln C_t$
Experiment
Prediction

#### Additional file 5 — Converged salt correction for duplex initialization.

Converged correction function fitted (left) to the difference $g_w^{\exp}(\rho) - g_w(\rho)$ of 18 duplexes data [2], The plot (right) of the predicted melting temperature correction versus the experiments of longer duplexes [3] shows a better agreement with Pearson correction $r = 0.54$.

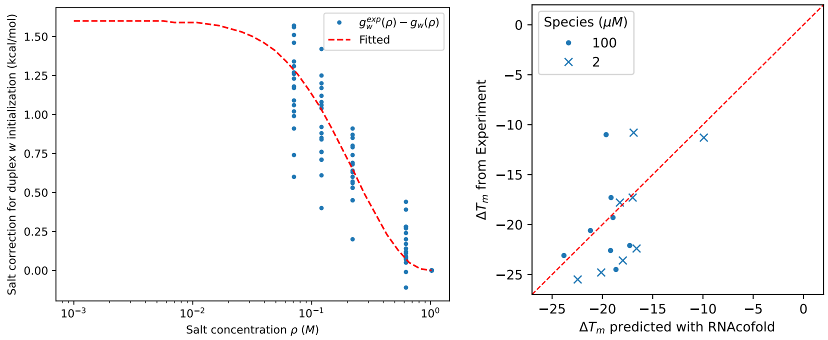


Converged salt correction for duplex initialization

$$g_{\text{init}}(\rho) = -\exp\left(a(\log(\frac{\rho}{\rho_0}))^2 + b\log(\frac{\rho}{\rho_0}) + \ln c\right) + c$$

with $a = -1.25480589$, $b = -0.05306256$, and $c = 160$. The parameter $c$ is a constant to ensure all data points are positive in natural logarithm while fitting.
